# Systematic analysis of the effects of the DNA damage response network in telomere defective budding yeast

**DOI:** 10.1101/101253

**Authors:** Eva-Maria Holstein, Greg Ngo, Conor Lawless, Peter Banks, Matthew Greetham, Darren Wilkinson, David Lydall

## Abstract

Functional telomeres are critically important to eukaryotic genetic stability. Budding yeast is a powerful model organism for genetic analysis and yeast telomeres are maintained by very similar mechanisms to human telomeres. Scores of proteins and pathways are known to affect telomere function. Here, we report a series of related genome-wide genetic interaction screens performed on budding yeast cells with acute or chronic telomere defects. We examined genetic interactions in cells defective in Cdc13 and Stn1, affecting two components of CST, a single stranded DNA (ssDNA) binding complex that binds telomeric DNA. We investigated genetic interactions in cells with defects in Rfa3, affecting the major ssDNA binding protein, RPA, which has overlapping functions with CST at telomeres. We also examined genetic interactions in cells lacking *EXO1* or *RAD9,* affecting different aspects of the DNA damage response in a *cdc13-1* background. Comparing fitness profiles across the data sets allows us build up a picture of the specific responses to different types of dysfunctional telomeres. Our results show that there is no universal response to telomere defects. To help others engage with the large volumes of data we make the data available via two interactive web-based tools: Profilyzer and DIXY. Among numerous genetic interactions we found the *chk1*Δ mutation improved fitness of *cdc13-1 exo1*Δ cells more than other checkpoint mutations (*ddc1*Δ, *rad9*Δ, *rad17*Δ, *rad24*Δ), whereas in *cdc13-1* cells the effects of all checkpoint mutations were similar. We find that Chk1 stimulates resection at defective telomeres, revealing a new role for Chk1 in the eukaryotic DNA damage response network.

## Introduction

The most important function of telomeres is to shield chromosome ends from being recognised as DNA double strand breaks (DSBs). The DNA damage response (DDR) to dysfunctional telomeres strongly affects genome stability, ageing and cancer (Gunes and Rudolph, 2013, Artandi and DePinho, 2010, Aubert and Lansdorp, 2008, Blackburn et al., 2015). In the eukaryotic model organism S. *cerevisiae,* the fitness of cells with defective telomeres can be increased, or decreased, by mutations affecting scores of different processes (Addinall et al., 2011). Analogous genetic interactions in human cells will presumably affect ageing and cancer.

Chromosome end protection depends on numerous proteins that bind both double stranded and single stranded telomeric DNA. In budding yeast, the CST complex, consisting of Cdc13, Stn1 and Ten1, binds telomeric single stranded DNA (ssDNA) and plays a critical role in telomere protection (Wellinger and Zakian, 2012). In budding yeast *CDC13*, *STN1* and *TEN1* are each essential genes (Price et al., 2010, Bertuch and Lundblad, 2006). The analogous genes in human and plant cells are CTC1, STN1 and TEN1 (Surovtseva et al., 2009, Chen et al., 2012). RPA is the major eukaryotic single-stranded DNA binding protein and plays critical roles in DNA repair, transcription and replication (Sugitani and Chazin, 2015). RPA is also critical for stimulating the DNA damage checkpoint pathway to cause cell cycle arrest. Interestingly, RPA, like CST, functions at telomeres; for example RPA binds telomeric ssDNA, promoting telomerase activity (Luciano et al., 2012). Analogously, CST has non-telomeric roles and was originally purified from human cells as "DNA Polymerase alpha accessory factor” (Goulian et al., 1990). Finally, there is evidence that different components of the CST complex perform different functions and that Cdc13, the largest sub-unit, is the least important component (Holstein et al., 2014, Lue et al., 2014, Lee et al., 2016). Therefore, there is much to learn about how the ssDNA binding proteins, CST and RPA, function at telomeres.

Inactivation of Cdc13 using the *cdc13-1* temperature sensitive allele leads to a complex and highly regulated DNA damage response. The DNA damage response results in extensive 5’-3’ telomeric DNA resection by two nuclease activities, Exo1 and Dna2-Sgs1 (Ngo and Lydall, 2010, Ngo et al., 2014). The ssDNA generated, which extends to single copy sub-telomeric loci, stimulates the DNA damage checkpoint kinase cascade, which phosphorylates many downstream targets to facilitate cell cycle arrest and DNA repair. The activation of this checkpoint response in *cdc13-1* strains is dependent on checkpoint sensors (the 9-1-1 complex, Ddc1, Mec3 and Rad17 in budding yeast), an adaptor (Rad9), a central kinase (Mec1), and effector kinases (Rad53 and Chk1). Checkpoint proteins also influence resection, notably the 9-1-1 complex stimulates resection while Rad9 and Rad53 inhibit resection (Jia et al., 2004, Zubko et al., 2004, Morin et al., 2008). Control over resection, just like control over cell cycle arrest, is critical for ensuring a measured response to telomere defects. This ensures that cells can recover when telomere defects are overcome.

To better understand the network that responds to telomere defects, we combined different types of defect with genome-wide libraries of mutations to create systematic double and triple mutant libraries. We then used quantitative fitness analysis (QFA) to measure fitness of these strains in response to chronic low level telomere defects or more acute telomere defects. We assessed fitness of strains with defects in Cdc13, Stn1 and Rfa3. In addition, we examined cells with telomere defects in combination with nuclease or checkpoint defects in *cdc13-1 exo1*Δ and *cdc13-1 rad9*Δ strains. Our results illustrate the complexity of the network that responds to telomere stress and clearly show that each telomere defect is different and that the pathways that respond to each defect are distinct. We followed up on a small subset of these interactions and discovered new roles for Chk1 in the response to uncapped telomeres. To help others interact with the large volumes of data we have made them available via two complementary web based interactive tools: DIXY and Profilyzer.

## Material and Methods

### Strains

All experiments were performed in W303 or S288C background strains (Table 1).

**Table 1.**
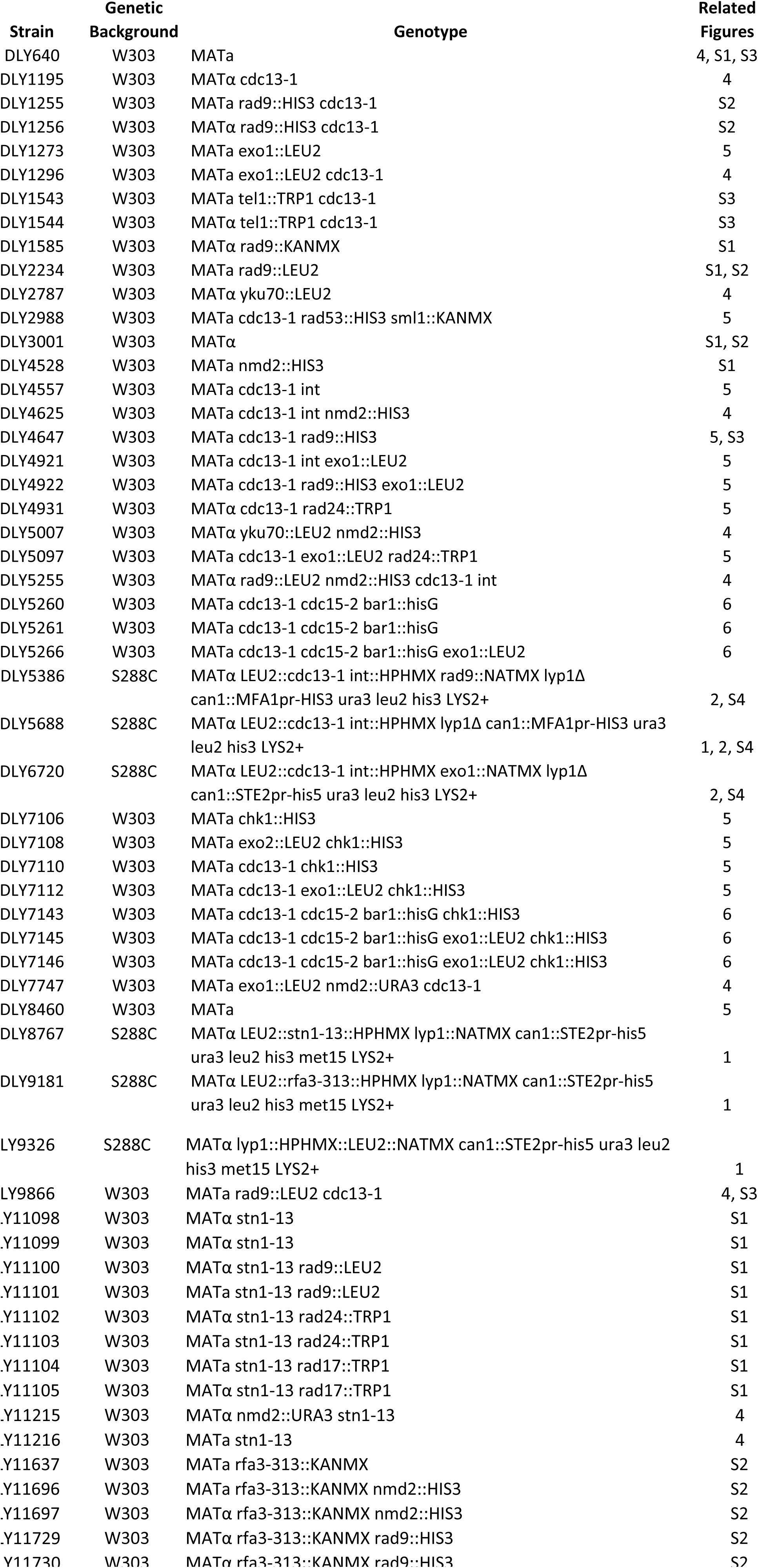

### QFA

Query strains used are described in Table 2. SGA (synthetic genetic array) was performed as previously described, crossing *cdc13-1*, *cdc13-1 rad9*Δ, *cdc13-1 exo1*Δ, *stn1-13*, *rfa3-313*, *lyp1*Δ, *ura3*Δ with the genome-wide single gene deletion knock out collection (Tong and Boone, 2006, Tong et al., 2001). *cdc13-1*, *stn1-13*, *rfa3-313* were flanked by the selectable HphMX and *LEU2* markers. Each strain also contains a third selectable marker, NATMX. In the *stn1-13, rfa3-313* query strains, NATMX is integrated at the *LYP1* locus, whereas in the *cdc13-1 rad9*Δ and *cdc13-1 exo1*Δ query strains, *RAD9* and *EXO1* were replaced by NATMX, respectively.

**Table 2.**
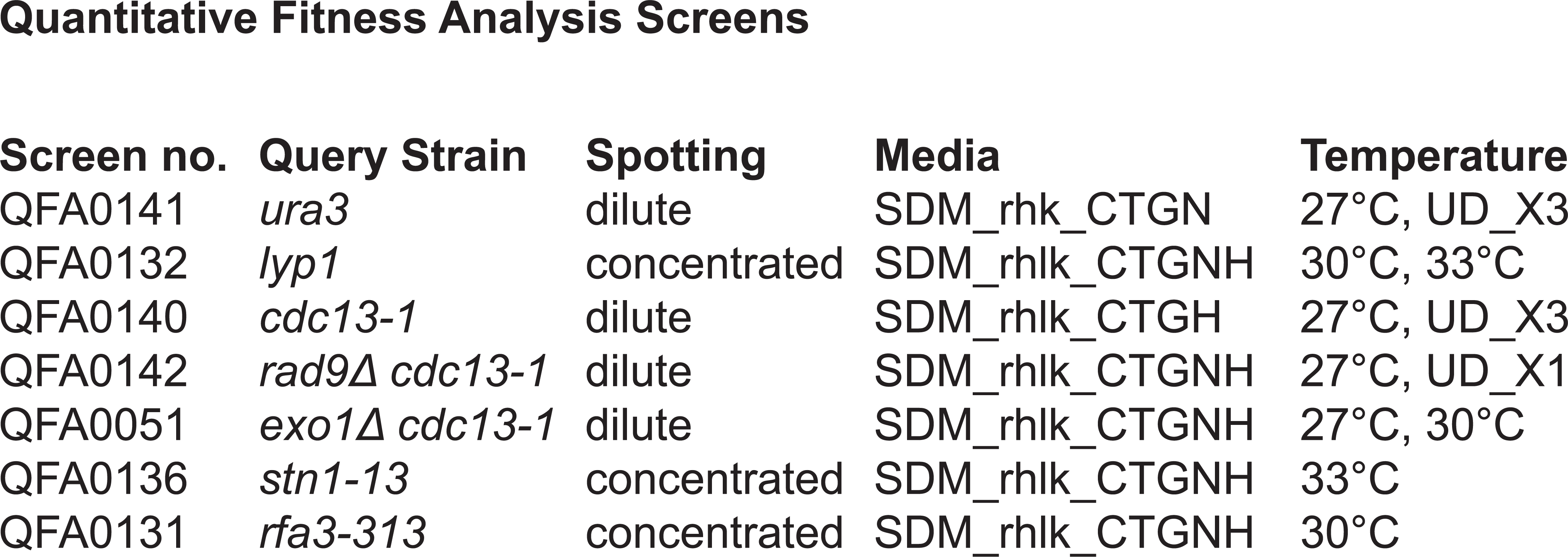
Quantitative Fitness Analysis Screens

For QFA strains were inoculated into 200 μl liquid CSM media in 96-well plates and grown for 2 days at 20°C without shaking. Saturated cultures were spotted onto solid agar plates either directly, or after diluting in water. Agar plates were incubated and imaged as before (Addinall et al., 2011, Dubarry et al., 2015). For the *ura3*Δ (UD) and the *cdc13-1* (UD) up-down assays, plates were incubated at 36°C, for 5h, followed by 20°C for 5h, three times, then plates were kept at 20°C for the remaining time. For the *rad9Δ cdc13-1* (UD) up-down assay, plates were incubated at 36°C for 8h, followed by incubation at 23°C for the remaining time.

### Profilyzer

Profilyzer is a web-based tool for visualizing and comparing results from multiple QFA screens at once (Dubarry et al., 2015). Profilyzer consists of various custom-built R functions inside a Shiny framework (Chang et al., 2015). A live instance of Profilyzer for this manuscript can be found on this web-page: http://research.ncl.ac.uk/qfa/Holstein2016. Fitness data and source code underlying this instance, as well as genetic interaction data underlying the instance of DIXY generated for this manuscript, can be found on GitHub: https://github.com/lwlss/Holstein2016.

### Small Scale Spot tests

Several colonies were inoculated into 2ml YEPD and incubated on a wheel at 23°C overnight until saturation. Five-fold serial dilutions of saturated cultures wer spotted onto agar plates using a 48 or 96-prong replica plating device. Plates were incubated at different temperatures for 2-3 days before being photographed.

### Cell cycle analysis

W303 strains containing *cdc13-1 cdc15-2 bar1*Δ mutations were grown at 23°C and arrested in G1 using alpha factor. Strains were then released from G1 at 36°C to induce telomere uncapping. Samples were taken periodically and cell cycle position was determined using DAPI staining (Zubko et al., 2004).

### QAOS

ssDNA levels were determined using Quantitative Amplification Of Single-stranded DNA (QAOS), as previously described (Holstein and Lydall, 2012).

## Results

Previous comparisons between genome-wide genetic interaction screens of telomere defective *cdc13-1* and *yku70*Δ strains revealed similarities and differences in the types of interactions observed (Addinall et al., 2011). We therefore wanted to extend this approach to other telomere-defective situations.

### STN1

We first examined genetic interactions affecting fitness of *stn1-13* mutants. Stn1, like Cdc13, is a component of the CST complex (Cdc13-Stn1-Ten1) that binds telomeric ssDNA. However, there is evidence that Cdc13 and Stn1 perform different functions. For example, a *stn1-186t* truncated allele is synthetically lethal in combination with the *rad9*Δ mutation (Petreaca et al., 2007), whereas the *cdc13-1* mutation is suppressed by *rad9*Δ (Zubko et al., 2004). This suggests that the molecular defects caused by *stn1-13* and *cdc13-1* mutations are different. Consistent with this hypothesis, mutations that completely bypass the requirement for *CDC13* and permit *cdc13*Δ cells to grow, do not bypass the requirement for *STN1* (Holstein et al., 2014).

A temperature sensitive *stn1-13* ts allele was crossed to a genome wide collection of mutations (*yfg*Δ) and fitness of resulting double mutants measured by QFA. *stn1-13* has a higher permissive temperature than *cdc13-1* and therefore we performed experiments at 33°C, a temperature that moderately inhibits growth of *stn1-13* strains. The overall pattern of genetic interactions observed in *stn1-13* cells is different to that previously reported for *cdc13-1* cells, with a tighter clustering of fitness measurements (Figures 1B, C). The different patterns could be due to the different properties of the two alleles, or because of technical differences in the two experiments (see Table 2), which were performed more than five years apart.

**Figure 1:**
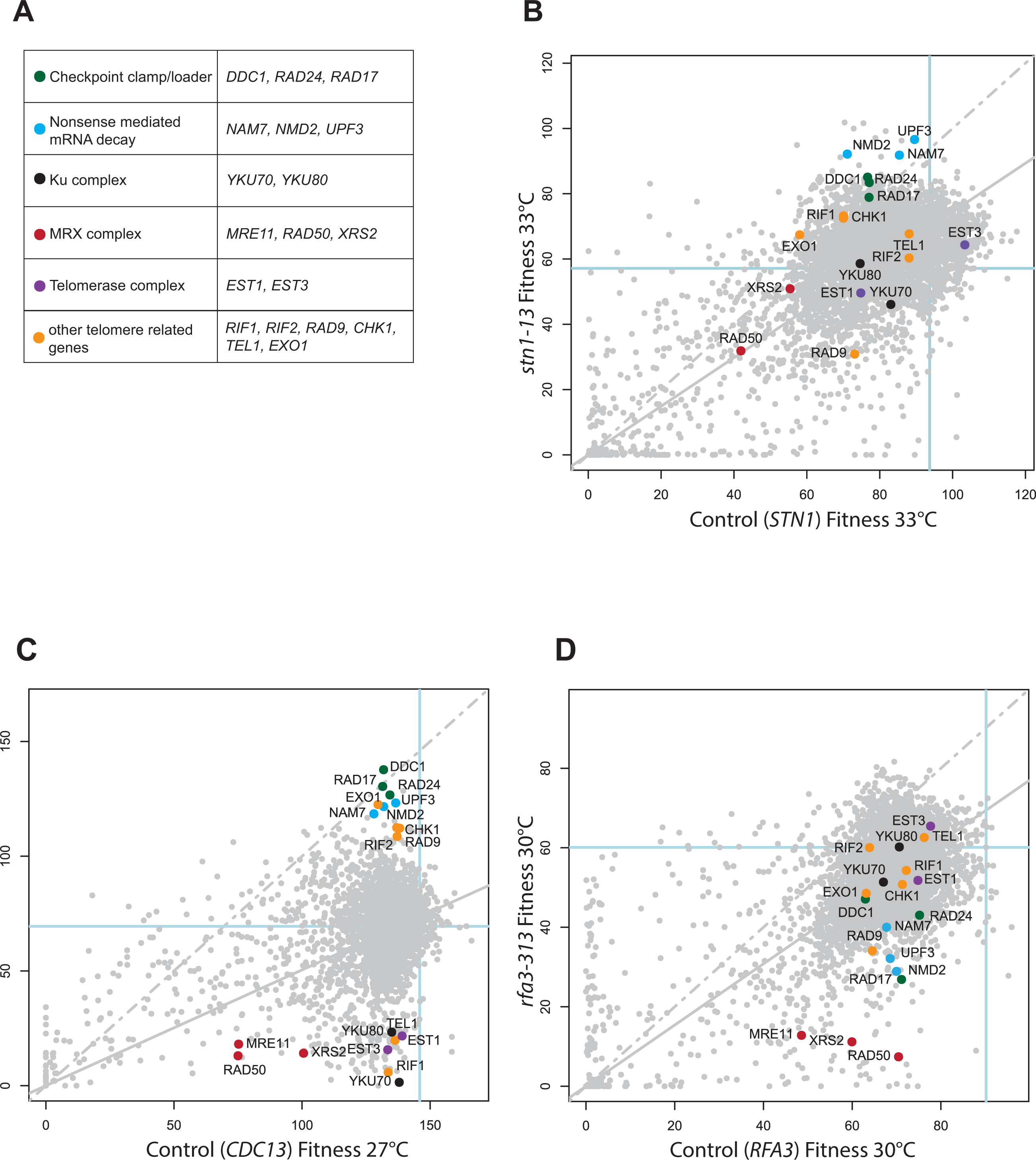
Genome-wide analysis of genetic interactions with mutations affecting the function of proteins that bind to ssDNA. A) Table of genes with functions at telomeres highlighted in B, C and D. B) Fitness plot showing evidence for genetic interaction between members of the yeast knockout collection and *stn1-13.* Each point summarises the effect of replicates of a library of *yfg*Δ mutations on *STN1 lyp1*Δ or *stn1-13* strain fitness at 33°C. Fitness is measured as MDR×MDP as previously described (Addinall et al., 2011). The dashed grey line represents the line of equal fitness in both strain backgrounds and solid grey is the predicted fitness assuming genetic independence. C) Same as in B but in *CDC13 ura3*Δ or *cdc13-1* backgrounds and at 27°C. D) Same as in B but in *RFA3 lyp1*Δ and *rfa3-313* contexts and at 30°C.

To assess the technical quality of the *stn1-13* experiment and to identify potentially informative genetic interactions, we highlight the positions of 19 diagnostic gene deletions known to play roles in telomere physiology or in telomere defective strains (Figure 1A and Table 3). In particular, among these 19, are five sets of related gene deletions affecting either the checkpoint sliding clamp, nonsense mediated mRNA decay, the Ku complex, the MRX complex or telomerase (Figure 1A). In principle, if members of a protein complex always function together and the complex function is dependent on all members, then each individual deletion affecting such a complex should have similar effects to other deletions and show very similar fitness patterns and genetic interactions. Therefore, assessing the reproducibility of the effects of deletions affecting the same complex is one way to assess the quality of the genetic interactions we have observed.

**Table 3.**
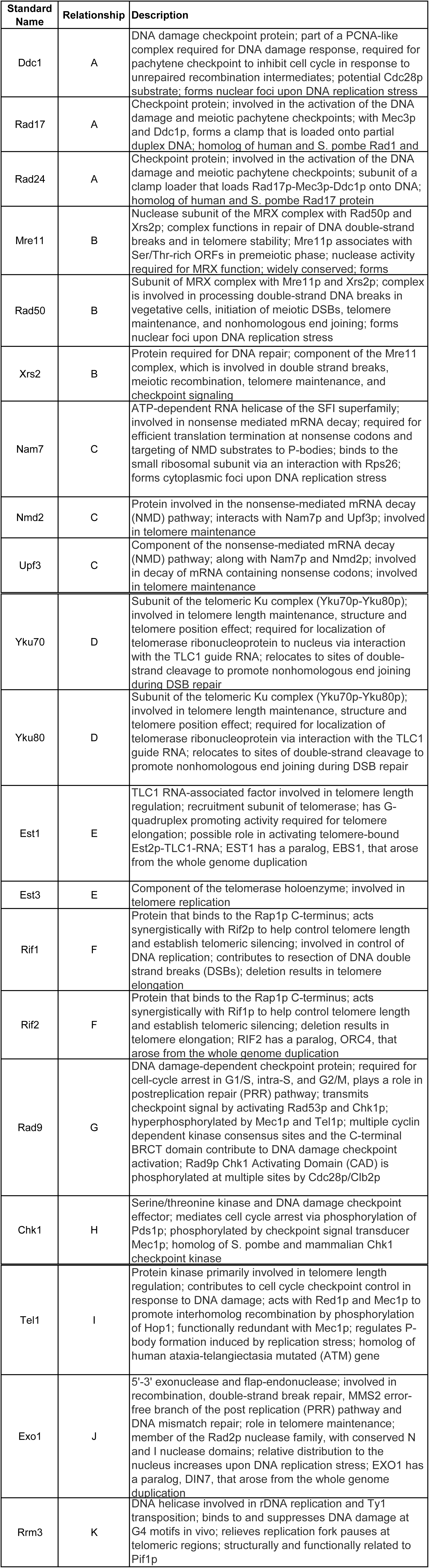
List of genes highlighted in the QFA screens

| Standard Name | Relationship | Description |
| --- | --- | --- |
| Ddc1 | A | DNA damage checkpoint protein; part of a PCNA-like complex required for DNA damage response, required for pachytene checkpoint to inhibit cell cycle in response to unrepaired recombination intermediates; potential Cdc28p substrate; forms nuclear foci upon DNA replication stress |
| Rad17 | A | Checkpoint protein; involved in the activation of the DNA damage and meiotic pachytene checkpoints; with Mec3p and Ddc1p, forms a clamp that is loaded onto partial duplex DNA; homolog of human and *S. pombe* Rad1 and |
| Rad24 | A | Checkpoint protein; involved in the activation of the DNA damage and meiotic pachytene checkpoints; subunit of a clamp loader that loads Rad17p-Mec3p-Ddc1p onto DNA; homolog of human and *S. pombe* Rad17 protein |
| Mre11 | B | Nuclease subunit of the MRX complex with Rad50p and Xrs2p; complex functions in repair of DNA double-strand breaks and in telomere stability; Mre11p associates with Ser/Thr-rich ORFs in premeiotic phase; nuclease activity required for MRX function; widely conserved; forms |
| Rad50 | B | Subunit of MRX complex with Mre11p and Xrs2p; complex is involved in processing double-strand DNA breaks in vegetative cells, initiation of meiotic DSBs, telomere maintenance, and nonhomologous end joining; forms nuclear foci upon DNA replication stress |
| Xrs2 | B | Protein required for DNA repair; component of the Mre11 complex, which is involved in double strand breaks, meiotic recombination, telomere maintenance, and checkpoint signaling |
| Nam7 | C | ATP-dependent RNA helicase of the SFI superfamily; involved in nonsense mediated mRNA decay; required for efficient translation termination at nonsense codons and targeting of NMD substrates to P-bodies; binds to the small ribosomal subunit via an interaction with Rps26; forms cytoplasmic foci upon DNA replication stress |
| Nmd2 | C | Protein involved in the nonsense-mediated mRNA decay (NMD) pathway; interacts with Nam7p and Upf3p; involved in telomere maintenance |
| Upf3 | C | Component of the nonsense-mediated mRNA decay (NMD) pathway; along with Nam7p and Nmd2p; involved in decay of mRNA containing nonsense codons; involved in telomere maintenance |
| Yku70 | D | Subunit of the telomeric Ku complex (Yku70p-Yku80p); involved in telomere length maintenance, structure and telomere position effect; required for localization of telomerase ribonucleoprotein to nucleus via interaction with the TLC1 guide RNA; relocates to sites of double-strand cleavage to promote nonhomologous end joining during DSB repair |
| Yku80 | D | Subunit of the telomeric Ku complex (Yku70p-Yku80p); involved in telomere length maintenance, structure and telomere position effect; required for localization of telomerase ribonucleoprotein via interaction with the TLC1 guide RNA; relocates to sites of double-strand cleavage to promote nonhomologous end joining during DSB repair |
| Est1 | E | TLC1 RNA-associated factor involved in telomere length regulation; recruitment subunit of telomerase; has G-quadruplex promoting activity required for telomere elongation; possible role in activating telomere-bound Est2p-TLC1-RNA; EST1 has a paralog, EBS1, that arose from the whole genome duplication |
| Est3 | E | Component of the telomerase holoenzyme; involved in telomere replication |
| Rif1 | F | Protein that binds to the Rap1p C-terminus; acts synergistically with Rif2p to help control telomere length and establish telomeric silencing; involved in control of DNA replication; contributes to resection of DNA double strand breaks (DSBs); deletion results in telomere elongation |
| Rif2 | F | Protein that binds to the Rap1p C-terminus; acts synergistically with Rif1p to help control telomere length and establish telomeric silencing; deletion results in telomere elongation; RIF2 has a paralog, ORC4, that arose from the whole genome duplication |
| Rad9 | G | DNA damage-dependent checkpoint protein; required for cell-cycle arrest in G1/S, intra-S, and G2/M, plays a role in postreplication repair (PRR) pathway; transmits checkpoint signal by activating Rad53p and Chk1p; hyperphosphorylated by Mec1p and Tel1p; multiple cyclin dependent kinase consensus sites and the C-terminal BRCT domain contribute to DNA damage checkpoint activation; Rad9p Chk1 Activating Domain (CAD) is phosphorylated at multiple sites by Cdc28p/Clb2p |
| Chk1 | H | Serine/threonine kinase and DNA damage checkpoint effector; mediates cell cycle arrest via phosphorylation of Pds1p; phosphorylated by checkpoint signal transducer Mec1p; homolog of S. pombe and mammalian Chk1 checkpoint kinase |
| Tel1 | I | Protein kinase primarily involved in telomere length regulation; contributes to cell cycle checkpoint control in response to DNA damage; acts with Red1p and Mec1p to promote interhomolog recombination by phosphorylation of Hop1; functionally redundant with Mec1p; regulates P-body formation induced by replication stress; homolog of human ataxia-telangiectasia mutated (ATM) gene |
| Exo1 | J | 5'-3' exonuclease and flap-endonuclease; involved in recombination, double-strand break repair, MMS2 error-free branch of the post replication (PRR) pathway and DNA mismatch repair; role in telomere maintenance; member of the Rad2p nuclease family, with conserved N and I nuclease domains; relative distribution to the nucleus increases upon DNA replication stress; EXO1 has a paralog, DIN7, that arose from the whole genome duplication |
| Rrm3 | K | DNA helicase involved in rDNA replication and Ty1 transposition; binds to and suppresses DNA damage at G4 motifs in vivo; relieves replication fork pauses at telomeric regions; structurally and functionally related to Pif1p |

Importantly, individual deletions affecting the checkpoint clamp/loader or nonsense mediated mRNA decay, caused similar increases in fitness of *stn1-13* strains (located near the top of Figure 1B). It is known that disabling the NMD pathway leads to overexpression of the CST components Ten1 and Stn1 (Dahlseid et al., 2003, Addinall et al., 2011) and an increase in Stn1 or Ten1 levels could explain the suppressing effect of NMD mutations in both *stn1-13* and *cdc13-1* backgrounds (Figures 1B,C). The suppressing effects of mutations affecting the checkpoint clamp/loader are most likely because telomere defects stimulate the DNA damage checkpoint pathway in *stn1-13* and *cdc13-1* cells. Interestingly, inactivation of the Ku complex (*yku70*Δ, *yku80*Δ), or telomerase (*est1*Δ, *est3*Δ) resulted in comparatively minor reduction of the fitness of *stn1-13* cells, in comparison with their effects on *cdc13-1* strain fitness (Figure 1B and C). Reassuringly, our high-throughput data reproduces the observation that fitness defects caused by *stn1* mutations are enhanced by *rad9*Δ (Figure 1B) (Petreaca et al., 2007), whereas the fitness defect of *cdc13-1* is suppressed by *rad9*Δ (*rad9*Δ is below the regression line in Figure 1B but above the regression line in Figure 1C) (Zubko et al., 2004). It is particularly interesting that *rad9*Δ enhances, whereas *rad17*Δ, *rad24*Δ and *ddc1*Δ mildly suppress, *stn1-13* fitness defects (Figure 1B). In contrast, *rad9*Δ, *rad17*Δ, *rad24*Δ and *ddc1*Δ each strongly suppress *cdc13-1* (Figure 1C). Importantly, we confirmed that *rad9Δ* enhances while *rad17*Δ and *rad24*Δ suppress *stn1-13* in small scale experiments in a different (W303) genetic background (Figure S1). Interestingly, it was previously observed that *rad9*Δ and *rad24*Δ mutations were synthetically lethal with a truncated *stn1-186t* allele, suggesting that checkpoint function is essential in cells with *stn1-186t* uncapped telomeres (Petreaca et al., 2007), but not *stn1-13* cells (this work). It will be interesting to understand the molecular basis of this difference between *stn1-186t* and *stn1-13.* Overall, it is clear that there are similarities and differences in the genetic interactions observed in *cdc13-1* and *stn1-13* strains, presumably reflecting the fact that two alleles cause similar, but distinct, telomere defects.

### RFA3

The heterotrimeric Replication Protein A (RPA), consisting of Rfa1, Rfa2 and Rfa3, binds ssDNA and plays critical roles in transcription, DNA replication and DNA repair. One view of the relationship between RPA and CST is that RPA binds ssDNA throughout the genome, including at telomeres, whereas CST specifically binds telomeric ssDNA (Gao et al., 2007). To better understand the relationship between CST and RPA we performed QFA on strains containing the temperature sensitive *rfa3-313* allele (Figure 1D).

Interestingly, deletions of members of the NMD complex (*nam7*Δ, *nmd2*Δ, *upf3*Δ) caused a decrease in fitness in *rfa3-313* mutants (Figure 1D), the opposite effect to what is observed in *stn1-13* and *cdc13-1* mutants (Figure 1B and 1C). Checkpoint mutations (*chk1*Δ, *rad9*Δ, *rad17*Δ, *rad24*Δ, *ddc1*Δ) slightly enhanced fitness defects or were comparatively neutral, in *rfa3-313* mutants. In a different genetic background we were able to confirm that *nmd*Δ mutations enhance whereas the *rad9*Δ checkpoint mutation is comparatively neutral in *rfa3-313* cells (Figure S2). Although we do not yet understand the molecular basis of these results, it is noteworthy that the pattern of genetic interactions observed in *rfa3-313* strains is markedly different to that seen in telomere-defective *cdc13-1* or *stn1-13* strains.

#### The effects of Rad9 and Exo1 on the response to *cdc13-1* defects

A large network of proteins coordinates the response of cells to damaged telomeres. Deletion of *RAD9*, a checkpoint gene, or *EXO1*, a nuclease gene, improves the fitness of *cdc13-1* strains grown at semi-permissive temperature (Figure 1C). But Rad9 and Exo1 contribute in distinct ways to fitness of *cdc13-1* mutants (Zubko et al., 2004). Rad9 is a critical component of the cell cycle arrest pathway that responds to *cdc13-1* defects and Rad9 also binds chromatin to inhibit nucleases that generate single stranded DNA in response to defective telomeres. Exo1 is one of the nucleases that generate ssDNA in *cdc13-1* and other telomere defective strains. Therefore, we were interested to screen the genome-wide knock out library for genes interacting with *exo1*Δ or *rad9Δ* in a *cdc13-1* background, to better define the structure of the DNA damage response network that is active in *cdc13-1* cells.

Figure 2 allows us to compare the effects of gene deletions in cells with the *cdc13-1* mutation with or without *exo1*Δ or *rad9*Δ mutations. The general effect of *rad9*Δ on the library of *cdc13-1 yfg*Δ mutants was to improve fitness, as seen by the increased fitness of most library strains. The global effects of *exo1*Δ are harder to discern because the fitness of *cdc13-1 exo1*Δ strains was measured at 30°C (to better assess the effects of the telomere defect).

**Figure 2:**
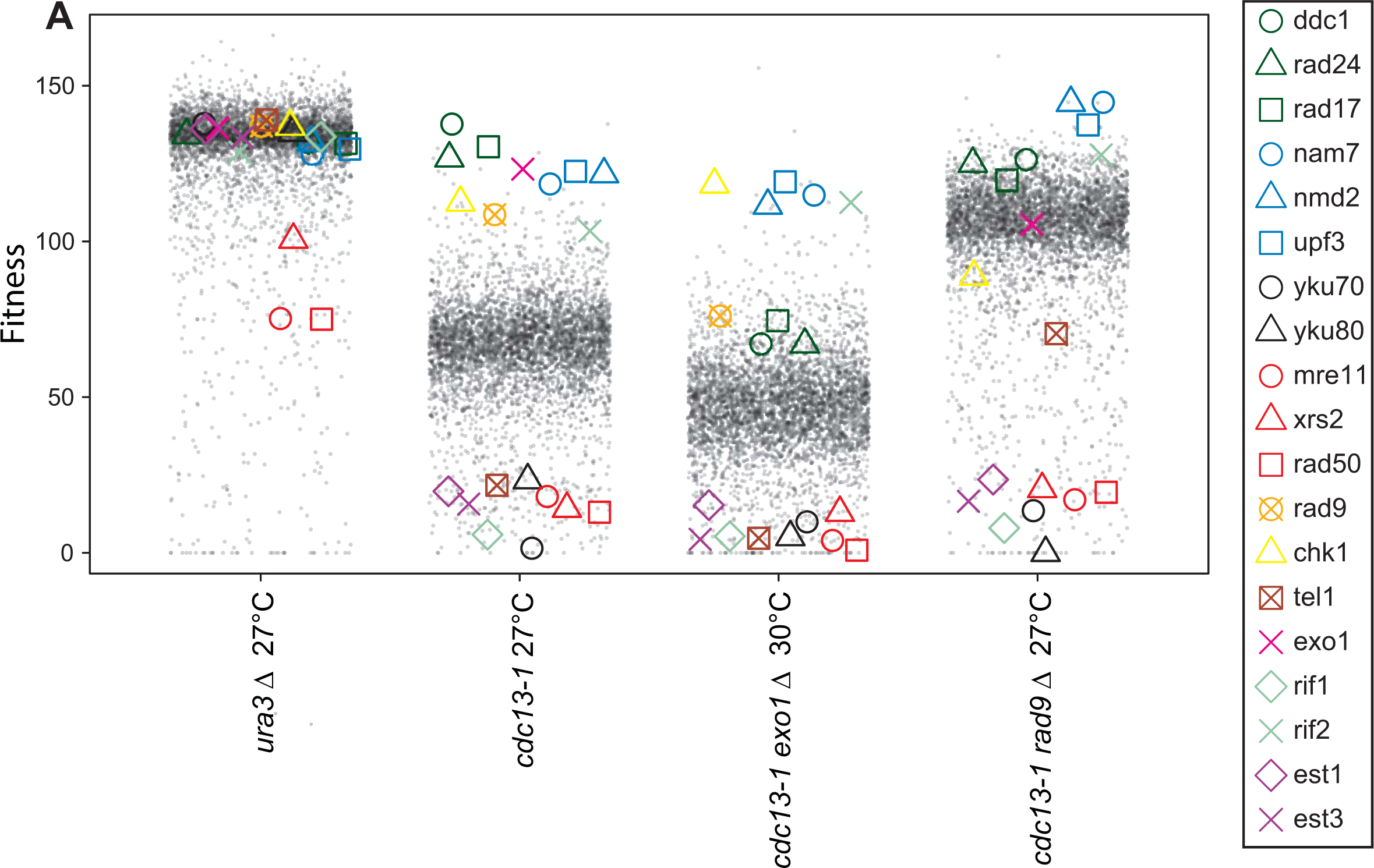
Effects of *rad9*Δ or *exo1*Δ on the fitness of *cdc13-1* strains. A) Fitness profile showing the effects of ~5,000 *yfg*Δ library mutations on the fitness of *ura3*Δ, *cdc13-1*, *cdc13-1 rad9*Δ and *cdc13-1 exo1*Δ strains. Each point represents the fitness of one gene deletion strain in each combination of genetic background and temperature. 19 telomere-related gene deletions are highlighted.

*cdc13-1 rad9*Δ (*yfg*Δ) strains are completely checkpoint defective while *cdc13-1 exo1*Δ cells are checkpoint proficient, but nuclease defective (Zubko 2004). It is interesting to compare different gene deletions in these contexts. For example, amongst the strongest suppressors of the *cdc13-1* fitness defect are deletions affecting the 9-1-1 complex (*rad17*Δ, *rad24*Δ, *ddc1*Δ), and *nmd*Δ mutations, affecting nonsense mediated decay (*nam7*Δ, *nmd2*Δ, *upf3*Δ). They show similar fitness in *cdc13-1* strains, but in *cdc13-1 rad9*Δ and *cdc13-1 exo1*Δ strains the *nmd*Δ mutations are clearly fitter than *911*Δ checkpoint mutations. It is notable that *chk1*Δ had a stronger effect than other checkpoint gene deletions in *cdc13-1 exo1*Δ strains and we investigate this further in Figure 6.

Numerous deletions affecting telomerase, the Ku complex, the MRX complex and Rif1, known to play important roles in telomere function, strongly reduced fitness in *cdc13-1*, *cdc13-1 rad9*Δ and *cdc13-1 exo1*Δ strains (Figure 2). *tel1*Δ showed a different pattern, it strongly reduced fitness in *cdc13-1* and *cdc13-1 exo1*Δ strains but less so in the *cdc13-1 rad9*Δ context (right column Figure 2). We were able to confirm the effect of *tel1Δ, rad9*Δ and both mutations in *cdc13-1* strains in small scale W303 spots tests (Figure S3).

#### Acute exposure to telomere defects

Growing *cdc13-1* cells at semi-permissive temperatures (e.g. 27°C) allows us to assess the effects of genes on fitness of cells with chronic, low level telomere defects and in this assay *RAD9* and *EXO1* have very similar effects (Figure 1C). A complementary approach is to identify those genes that affect the viability of *cdc13-1* mutants exposed to acute, high-level damage (Addinall et al, 2008). Using this approach, *RAD9* and *EXO1* have opposite effects. Exo1 reduces viability while Rad9 protects viability of *cdc13-1* cells (Zubko et al., 2004). The different effects of *RAD9, EXO1* and other genes can be rationalized by their effects on ssDNA accumulation at uncapped telomeres, with Exo1 stimulating ssDNA production and Rad9 inhibiting production (Zubko et al., 2004, Jia et al., 2004).

To identify genes that affect cell fitness after acute exposure to telomere defects we performed genome-wide experiments in which cells were exposed to acute periods of incubation at 36°C followed by recovery at 23°C. We call this type of temperature cycling protocol an up-down assay (UD). Importantly, the previously reported opposing effects of *rad9*Δ and *exo1*Δ on viability of *cdc13-1* cells in up-down assays were confirmed in our genome-wide experiments, with *exo1*Δ strains being amongst the fittest strains in the *cdc13-1* (UD) context and *rad9*Δ strains being amongst the least fit (Figure 3A) (Zubko et al., 2004, Addinall et al., 2008). In other contexts, *ura3*Δ (27°C), *ura3*Δ (UD) and *cdc13-1* (27°C), *exo1*Δ and *rad9*Δ had similar effects. We also observed that *rad17*Δ, *ddc1*Δ and *rad24*Δ checkpoint defective strains were less sensitive than *rad9Δ* strains in the *cdc13-1* (UD) context, but still somewhat sensitive, as has been reported before for *rad24*Δ and is discussed below (Figure 3A) (Zubko et al., 2004).

**Figure 3:**
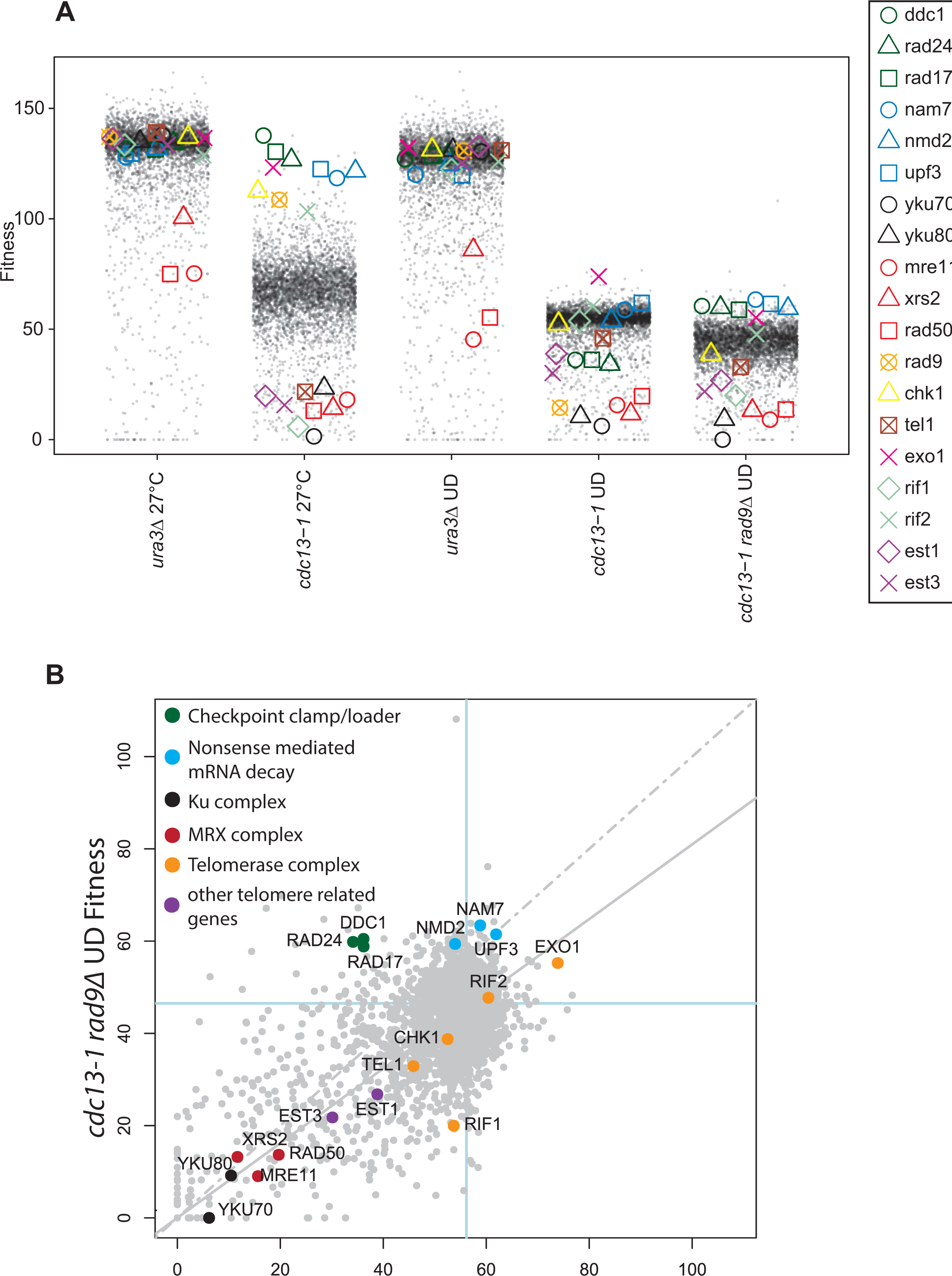
The effects of a library of *yfg*Δ mutations on fitness of cells after exposure to chronic or acute telomere defects. A) Fitness profile comparing the effects of ~ 5,000 *yfg*Δ library mutations on the fitness of strains indicated after chronic (27°C) or acute (UD) exposure to telomere defects. In UD (Up-Down) experiments, cells were exposed to short periods of incubation at 36°C (see methods). Data are plotted as in Fig 2A. B) Fitness plot comparing evidence for genetic interactions between *rad9*Δ and ~ 5,000 *yfg*Δ deletions in a *cdc13-1* background after acute telomere uncapping. Data are plotted as in Figure 1.

We observed in UD assays that *yku70*Δ and *yku80*Δ mutations reduced fitness of *cdc13-1* and *cdc13-1 rad9*Δ cells more than *est1Δ* and *est3Δ* mutations, affecting telomerase (Figure 3A). This pattern contrasts to what was seen at 27°C, after chronic low level *cdc13-1* telomere damage, when the effects of *yku70Δ, yku80Δ, est1*Δ and *est3*Δ were all similar to each other. The effects of the Ku heterodimer at high temperature in *cdc13-1* strains observations may be because the Ku heterodimer protects telomeres from the Exo1 nuclease, particularly at high temperature (Maringele and Lydall, 2002) and because Ku and Cdc13 function redundantly to cap the telomere (Polotnianka et al., 1998).

It is known that the 9-1-1 complex and Exo1 contribute to ssDNA production and cell death of *rad9*Δ *cdc13-1* strains but whether other proteins behave similarly, or have the opposite effect and maintain viability is unknown (Zubko et al., 2004). Most gene deletions that affected viability of *cdc13-1 rad9*Δ strains showed similar effects in *cdc13-1* strains and lie along the regression line in (Figure 3B). This was true for both suppressors of *cdc13-1*, like *exo1*Δ, and enhancers of *cdc13-1*, like *yku70*Δ and *yku80*Δ. However, there were clearly some gene deletions that behaved differently in *cdc13-1 rad9*Δ versus *cdc13-1* strains. *rif1*Δ, made cells considerably sicker in the *rad9Δ cdc13-1* (UD) context. Conversely, the *nmd*Δ mutations improved *rad9Δ cdc13-1* (UD) fitness more than *cdc13-1* (UD) fitness, suggesting that NMD has stronger effects on ssDNA accumulation in the absence or Rad9 (Holstein et al., 2014). It was notable that mutations affecting the checkpoint sliding clamp (*ddc1*Δ, *rad24*Δ and *rad17*Δ) clustered very tightly in Figure 3B. All strongly improved fitness in the checkpoint-defective *cdc13-1 rad9*Δ (UD) context, while reducing fitness in the *cdc13-1* (UD) context. This can be explained by the fact that *ddc1*Δ, *rad24*Δ and *rad17*Δ are each required for checkpoint arrest of *cdc13-1 RAD9* cells and contribute to rapid ssDNA production in *cdc13-1 rad9*Δ cells at 36°^C^ (Ngo et al., 2014, Booth et al., 2001). Overall, then, there are many informative differences in the effects of gene deletions of the viability of *cdc13-1* and *cdc13-1 rad9*Δ strains after acute exposure to telomere defects.

#### Effects of different gene deletions across several telomere defective strains

Comparison of the genome wide data sets shown in Figures 1-3 potentially reveal thousands of genetic interactions that are informative about telomere biology. Out of necessity, we have only discussed a tiny fraction of these. Therefore, to allow others to interact with these data, to identify other informative genetic interactions, we have also placed all our data into two web tools that facilitate data interrogation (Dubarry et al., 2015). The first tool DIXY (Dynamic Interactive XY plots), permits interaction with fitness data in a format similar to Figure 1. DIXY also allows regular scatter plots without interpretation in terms of genetic interaction to be generated and for any gene or genes to be highlighted across the plots. For example, Figure S4 shows a number of pairwise comparisons of the data in Figure 2, including fitness plots and regular scatter plots. DIXY allows users to readily monitor the changing positions of selected gene deletions across several plots.

In addition, all of the screens discussed here are accessible via Profilyzer (http://research.ncl.ac.uk/qfa/Holstein2016) a tool which allows examination of the effects of mutations across all the screens and which generates interactive online plots similar to Figure 2. Figure 4A illustrates the use of Profilyzer and shows very similar fitness profiles of three gene deletions affecting nonsense-mediated mRNA decay complex (*nam7*Δ, *nmd2*Δ and *upf3*Δ mutations (abbreviated to *nmd*Δ)) across all the screens. The NMD genes had minor effects on fitness of control strains (*ura3*Δ, *lyp1*Δ), increased fitness of *cdc13-1, stn1-13, cdc13-1 exo1*Δ, *cdc13-1 rad9*Δ strains, and *rad9Δ cdc13-1* strains after up-down treatments. In contrast, the *nmd*Δ mutations exacerbated fitness defects of *yku70*Δ and *rfa3-313* strains. Figure 4B confirms that the *nmd2*Δ mutation recapitulates many of these interactions in the different W303 genetic background, in low throughput spot tests on rich media (YEPD). We have also previously reported that *nmd2*Δ *rad9*Δ *cdc13-1* and *nmd2*Δ *cdc13-1* strains are comparatively viable in small scale up-down assays and the effect of *nmd2Δ* correlates with lower levels of ssDNA production at defective telomeres in *nmd2Δ* mutants (Holstein et al., 2014).

**Figure 4:**
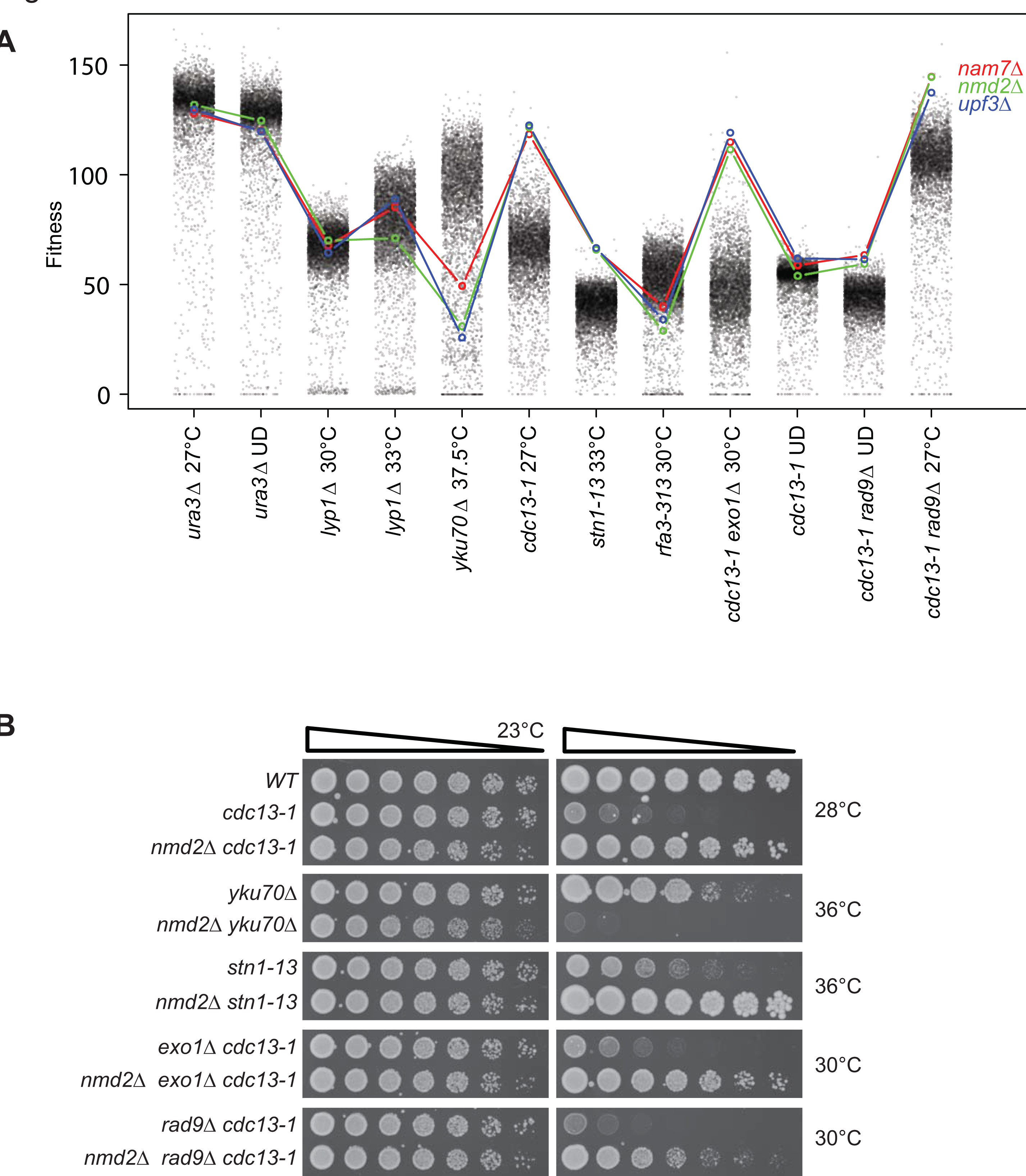
Effects of gene deletions affecting the Nonsense-Mediated mRNA Decay pathway across a range of telomere defective backgrounds. A) Profilyzer fitness profiles comparing the effects of *nam7*Δ, *nmd2*Δ, *upf3*Δ mutations on fitness across all the genome-wide screens presented in Figures 1-3. B) Saturated cultures of the yeast strains indicated (see Table 1) were five-fold serially diluted in water, spotted onto YEPD agar plates and incubated at the indicated temperatures for two days before being photographed.

Profilyzer was also used to compare the fitness profiles of *chk1*Δ, *ddc1*Δ, *rad9*Δ, *rad17*Δ and *rad24*Δ, five deletions with well-established effects on the response to uncapped telomeres (Figure 5A). In addition to comparing fitness profiles of individual genes or complexes, Profilyzer permits identification of those gene deletions with fitness profiles most similar to a query gene deletion across some or all screens. To illustrate this, we identified the 11 most similar profiles to *rad17*Δ. Reassuringly, given their known functions, *rad24*Δ and *ddc1*Δ were the closest gene deletions to *rad17*Δ based on their profile (Figure 5B). Other checkpoint mutations, *rad9*Δ and *chk1*Δ, also shown in Figure 5A, were also in the top 100 (out of 5,000) similar profiles (Figure 5B), reflecting their similar but distinct profiles to *rad17*Δ *(rad24Δ and ddc1Δ)*.

**Figure 5:**
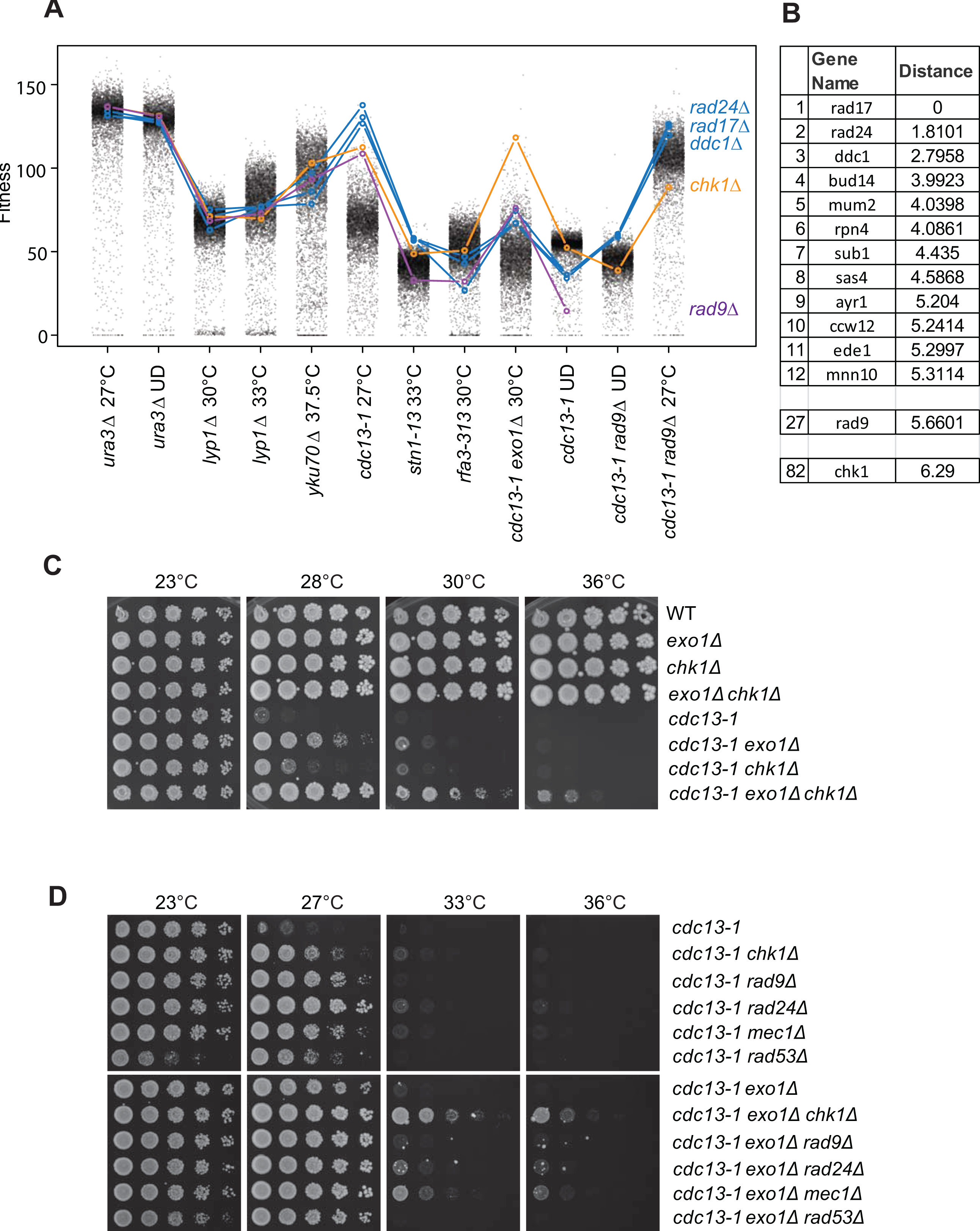
Effects of gene deletions affecting the DNA damage checkpoint pathway across a range of telomere defective backgrounds. A) Profilyzer fitness profiles comparing the effects of *ddc1*Δ, *rad24*Δ, *rad17*Δ, *rad9*Δ, *chk1*Δ mutations on fitness across all the screens presented in Figures 1-3. B) List of some of the gene deletions with most similar fitness profiles to *rad17*Δ out of ˜5,000 examined, including *rad9*Δ (position 27) and *chk1*Δ (position 82). C-D) Yeast cultures treated as in Figure 4B.

Rad17, Ddc1 (and Mec3) encode the 9-1-1 complex that is loaded at uncapped telomeres by the Rad24 sliding clamp loader and stimulates resection (Ngo et al., 2014). Rad9 binds chromatin near uncapped telomeres and inhibits resection. Chk1 is a kinase with a partial role signaling arrest of *cdc13-1* strains, while Ddc1, Rad9, Rad17 and Rad24 are essential for cell cycle arrest of *cdc13-1* strains (Lydall and Weinert, 1997). Figure 5A shows that in most contexts the effects of all five gene deletions were similar, as expected given their role in the DNA damage checkpoint pathways. For example, *chk1*Δ, *ddc1*Δ, *rad9*Δ, *rad17*Δ and *rad24*Δ all suppressed fitness defects when combined with *cdc13-1* and *exo1*Δ *cdc13-1* (Figure 5A). However, *rad9*Δ and/or *chk1*Δ behave differently to 9-1-1 complex mutations particularly in the context of *cdc13-1*, *stn1-13*, *cdc13-1 exo1*Δ or *cdc13-1 rad9*Δ mutations. In comparison with *ddc1*Δ, *rad9*Δ, *rad17*Δ and *rad24*Δ, *chk1*Δ had a stronger suppressive effect in *exo1*Δ *cdc13-1* strains, an observation reproduced in small scale spot tests in W303 (Figure 5C). Another striking difference in phenotypes was between *rad9*Δ and *rad17*Δ, *ddc1*Δ and *rad24*Δ in *stn1-13* mutants, *rad9Δ* enhanced fitness defects, while *rad17Δ, ddc1*Δ and *rad24*Δ suppressed the fitness defects (Fig 5A, Fig 1 and discussed earlier). All checkpoint mutations enhanced fitness defects in the *cdc13-1* up-down assay (Figure 5A), presumably because the loss of DNA damage checkpoint permits cell division in the context of telomere degradation, resulting in loss of viability.

#### Chk1 affects ssDNA production

*chk1*Δ was amongst strongest suppressors in the *cdc13-1 exo1*Δ context and sometimes in a separate location from other non-essential DNA damage checkpoint genes (*ddc1*Δ, *rad9*Δ, *rad17*Δ and *rad24*Δ) in other contexts (Figure 5A). To confirm the effects of Chk1 we performed a low throughput experiment in another genetic background (W303). We found that both *exo1*Δ and *chk1*Δ suppressed the temperature sensitivity of *cdc13-1* strains at 28°C and that double mutations strongly suppressed the temperature sensitivity of *cdc13-1* strains, permitting some growth at 36°C (Figure 5C).

To better understand the interactions between *exo1*Δ and *chk1*Δ in cells with uncapped telomeres, we combined *exo1*Δ with other checkpoint mutations in the same *cdc13-1* genetic background. Interestingly, *chk1*Δ was a stronger suppressor than *rad9*Δ, *rad24*Δ, *mec1*Δ or *rad53*Δ, suggesting that the suppressing effect of Chk1 may be independent of checkpoint activation and/or cell cycle arrest (Figure 5D). We conclude that *chk1*Δ is a particularly strong suppressor of *cdc13-1 exo1*Δ growth defects and this is most likely due to Chk1 having a checkpoint-independent role or roles.

Exo1 affects DNA resection at uncapped telomeres and DSBs, working in parallel to Dna2/Sgs1. Therefore, we hypothesized that Chk1, like Exo1, may stimulate resection at uncapped telomeres. To test this hypothesis, we examined resection at telomeres in synchronous cultures of *cdc13-1* strains at high temperature. We quantified the product of resection, ssDNA, by using Quantitative amplification of single-stranded DNA (QAOS) assay (Booth et al., 2001).

We examined ssDNA at two loci, *Y’600* and *Y’5000,* located in the Y’ subtelomeric elements, which are present in two thirds of budding yeast chromosome ends (including right telomere of chromosome V) (Figure 6A). In addition, we quantified ssDNA accumulation at a single copy locus *YER186C*, located 15kb from the right arm telomere of chromosomes V (Figure 6A). We first arrested *cdc13-1* cells in G1 with alpha-factor at 23°C, washed off the alpha factor, and incubated the cells at 36°C during the course of the experiment to induce telomere uncapping. To ensure a single round of DNA replication in all strains, we also utilized the *cdc15-2* allele to keep any checkpoint-deficient strains in late anaphase (Lydall and Weinert, 1995). Consistent with previous findings, we detected accumulation of 3’ ssDNA at *Y’600* and *Y’5000* in the wild type strains after one hour and further away at *YER186C* after two hours (Figure 6B) (Zubko et al., 2004). Importantly, we detected lower levels of ssDNA in *chk1Δ* mutants at all loci examined, suggesting that Chk1 does indeed stimulate telomere resection. The effect of *chk1*Δ was not as strong as *exo1*Δ, where ssDNA levels were very low, consistent with previous data (Zubko et al., 2004). Interestingly deleting *CHK1* in *cdc13-1 exo1*Δ mutants slightly reduced ssDNA, especially at Y’600, suggesting that Chk1 may stimulate Sgs1 dependent resection. However, the effect of Chk1 on resection is small, most likely because Sgs1-dependent resection is weak in *cdc13-1* or *cdc13-1 exo1Δ* strains (Ngo et al., 2014, Ngo and Lydall, 2010).

**Figure 6:**
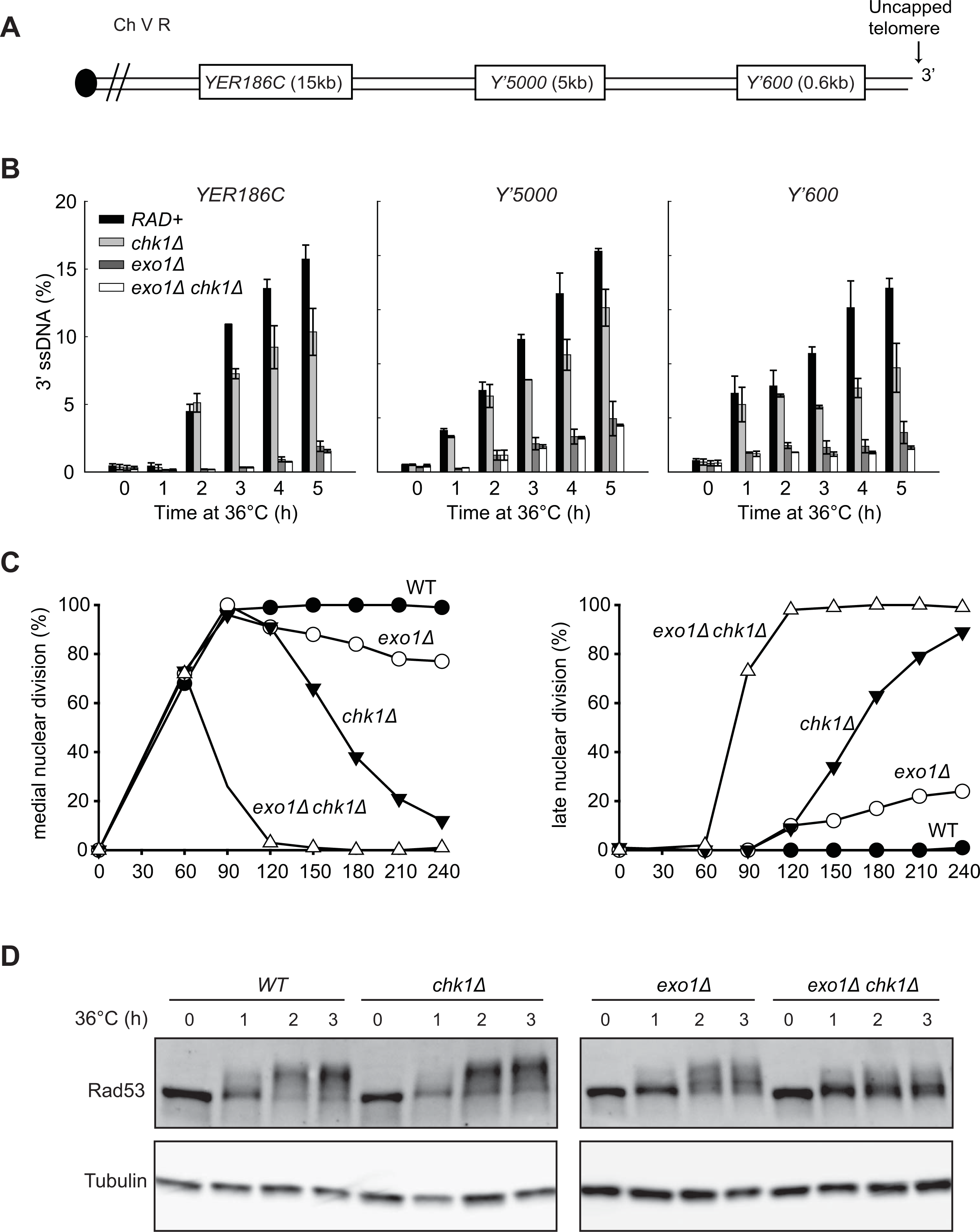
Chk1 stimulates resection, Rad53 phosphorylation and checkpoint activation in response to telomere defects. A) Map of the right arm of Chromosome V. B) Quantification of 3’ ssDNA accumulation at loci indicated following telomere uncapping. All strains contain *cdc13-1 cdc15-2 bar1*Δ mutations (Table 1). The data and error bars plotted are means and standard error of the mean from two independent experiments. C) Cell cycle position of the indicated *cdc13-1 cdc15-2 bar1Δ* strains was assessed by counting DAPI stained cells and the percentage of cells at medial or late nuclear was determined. D) Yeast strains with the indicated genotypes (all in *cdc13-1 cdc15-2 bar1*Δ background) were subjected to western blot analysis with anti-Rad53 and anti-tubulin antibodies following telomere uncapping.

To further establish the role of Chk1 in Sgs1-dependent resection, we examined Rad53 phosphorylation as a marker for DNA resection. We previously showed that Sgs1 stimulates Exo1-independent Rad53 phosphorylation in *cdc13-1 exo1*Δ strains (Ngo and Lydall, 2010). To test whether Chk1 stimulates resection by Sgs1, we examined whether Chk1 stimulates Rad53 phosphorylation in *cdc13-1 exo1*Δ strains (Figure 6C). Consistent with previous findings, we detected Rad53 phosphorylation in *cdc13-1* strains after 2 hours at 36°C and this phosphorylation was slightly redcued in *cdc13-1 exo1*Δ strains (Figure 6C) (Ngo and Lydall, 2010). Like *sgs1*Δ, *chk1*Δ did not reduce Rad53 phosphorylation in *cdc13-1* strains (Figure 6C) (Ngo and Lydall, 2010). However, *chk1*Δ ( like *sgs1*Δ), reduced Rad53 phosphorylation in *cdc13-1* exo1Δ strains, showing that Chk1 stimulates Sgs1-dependent resection and Rad53 phosphorylation (Ngo and Lydall, 2010).

Our previous study shows that inactivating both Exo1 and Sgs1 pathways of resection was not enough to permit *cdc13-1 exo1*Δ *sgs1*Δ cells to grow at 36°C, because Rad9-dependent cell cycle arrest can still be activated (Ngo and Lydall, 2010). So why do *cdc13-1 exo1*Δ *chk1*Δ cells grow at 36°C (Figure 5C)? We hypothesized that this is because in addition to inactivating Sgs1-dependent resection and Rad53 phosphorylation, *chk1*Δ (by inactivating Pds1) also strongly inactivates cell cycle arrest in *cdc13-1 exo1*Δ strains (Sanchez et al., 1999). To test this hypothesis, we examined cell cycle arrest of *cdc13-1, cdc13-1 chk1*Δ, *cdc13-1 exo1*Δ and *cdc13-1 exo1*Δ *chk1*Δ strains, assessing the fraction of cells arrested by *cdc13-1*, at medial nuclear division. Consistent with previous findings, *cdc13-1* cells remained arrested for at least 4 hours (Figure 6C) (Zubko et al., 2004). Both *exo1*Δ and *chk1*Δ strains showed mild checkpoint defects as small fractions (about 10%) of *exo1*Δ or *chk1*Δ cells failed to maintain arrest at the medial nuclear division stage at 2 hours. At later times *chk1*Δ strains showed a more severe checkpoint defect than *exo1Δ* strains, such that by four hours more than 80% of *chk1*Δ cells escaped arrest in comparison to 20% of *cdc13-1 exo1*Δ. *chk1*Δ caused a strong defect in checkpoint activation in *exo1Δ* background, as essentially all *exo1Δ chk1Δ* cells failed to arrest at medial nuclear division. We conclude that Chk1 stimulates Exo1-independent DNA resection and DNA damage checkpoint activation in *cdc13-1* strains and the inactivation of DNA resection and DNA checkpoint activities permits *cdc13-1 exo1*Δ *chk1*Δ cells to grow at 36°C.

## Discussion

Yeast telomeres are in many ways very similar to human telomeres, most notably relying on telomerase as a means to overcome the end replication problem. In this paper we have systematically explored genetic interactions that suppress or enhance different types of telomere defect, or affect fitness of cells with telomere defects in combination with mutations affecting aspects of the DNA damage response. These new data extend previous analyses of telomere defective *cdc13-1* and *yku70*Δ yeast strains (Addinall et al., 2011). We have also analysed how genes affect fitness after either acute or chronic telomere exposure, mimicking different classes of telomere defect likely to arise in human cells. For comparison, we also examined genetic interactions with a mutation affecting RPA, the central ssDNA binding protein, with roles in general chromosome and telomere replication and repair.

Our experiments clearly show that each telomere defect shows distinct genetic interactions, with only partially conserved, genetic suppressor and enhancer interactions. Overall then, it is clear that there is no universal response to telomere defects and therefore there is no single mechanism to overcome the adverse effects of telomere dysfunction. These observations in yeast are consistent with data from humans showing that mutations affecting telomere maintenance proteins cause different diseases. Individuals inheriting identical mutations can present with variable symptoms, or no symptoms, presumably, at least in part, because other inherited mutations suppress or enhance phenotypes (Armanios et al., 2007, Holohan et al., 2014).

Our work also clearly illustrates that a complex network of interactions responds to telomere defects and that inactivation of genes that play important roles in this network (e.g. *RAD9* and *EXO1*), changes the effects of other genes in the network. Consistent with this we recently reported that *cdc13-1* mutants lacking *RAD9* or *EXO1* retain the ability to adapt to low level telomere damage (Markiewicz-Potoczny and Lydall, 2016).

We saw many interesting patterns across the genome-wide datasets. The behaviour of *exo1*Δ and *nmd*Δ mutations are particularly interesting. These mutations seem to suppress most telomere defects, but are comparatively neutral, or enhance, the *rfa3-313* fitness defect, affecting general DNA replication. We have previously shown that *exo1*Δ and *nmd*Δ mutations reduce ssDNA levels near uncapped telomeres (Holstein et al., 2014). This mechanism most likely explains why *exo1*Δ and *nmd*Δ mutations suppress both chronic and acute telomere defects. Although, we note that *nmd*Δ mutations exacerbate fitness defects of *yku70*Δ mutants, at high temperature, which is also associated with increased telomeric ssDNA (Addinall et al., 2011, Maringele and Lydall, 2002).

It is interesting that *rad9*Δ, a checkpoint mutation affecting the yeast homologue of human 53BP1, suppresses *cdc13-1* telomere defective mutants growing with chronic telomere defects, but enhances fitness defects in nearly every other situation we tested, including *cdc13-1* strains exposed to acute telomere defects and *stn1-13* and *rfa-313* cells growing with chronic defects. Other checkpoint mutations, most clearly *rad17*Δ, *rad24*Δ and *ddc1*Δ, showed different patterns. We strongly suspect that the different effects of Rad9 in the different telomere defective contexts are due to its dual roles, inhibiting ssDNA accumulation and signalling cell cycle arrest. Rad9 binds chromatin and inhibits ssDNA accumulation (Lazzaro et al., 2008). The strong suppression of *cdc13-1* by *exo1*Δ *chk1*Δ double mutations may be explained by the novel finding that Chk1 contributes to ssDNA production in *cdc13-1* cells, a new role for Chk1 in the DNA damage response network.

It is clear that *cdc13-1* and *stn1-13,* affecting two components of the CST complex, show very different genetic interactions. At face value these differences are inconsistent with the idea that the CST complex functions as a single entity. Indeed, our favoured explanation for these data is that Stn1 performs different functions to Cdc13. We have previously shown that genetic interactions that suppress *cdc13*Δ do not suppress *stn1*Δ (Holstein et al., 2014). There is also biochemical evidence that Stn1 or Ten1 facilitates DNA replication without help from Cdc13 (Lue et al., 2014) and acts as a molecular chaperone in plants (Lee et al., 2016). It is also possible that each allele does not cause gradual loss of function of Cdc13 or Stn1, but rather causes separations of function. We think this is unlikely, in part because *cdc13-1* cells grown at high temperature behave like *cdc13*Δ cells (Garvik et al., 1995), but further experiments will be necessary to distinguish between models.

The volume of work we report in this paper is somewhat overwhelming. The experiments are informative and potentially of interest to many interested in telomere biology, DNA replication and chromosome biology. We have only been able to follow up and confirm a very small number of the interactions we observed in high throughput experiments. However, to help others interrogate the data we have uploaded all our data into two interactive web tools that allow users to interact with and search the data in various ways. We hope that our data will help others gain new insights into telomere biology.

## Acknowledgements

We thank Simon Cockell for help with Dixy and Adrian Blackburn for help with robotic analysis. We thank the MRC (MR/L001284/1), BBSRC (BB/M002314/1) and Wellcome Trust for support (WT093088MA).

---

**Table S1: Strains used in this study**

**Table S2: Query strains used for QFA**

**Table S3: Genes with functions at telomeres highlighted throughout this study**

**Figure S1:**
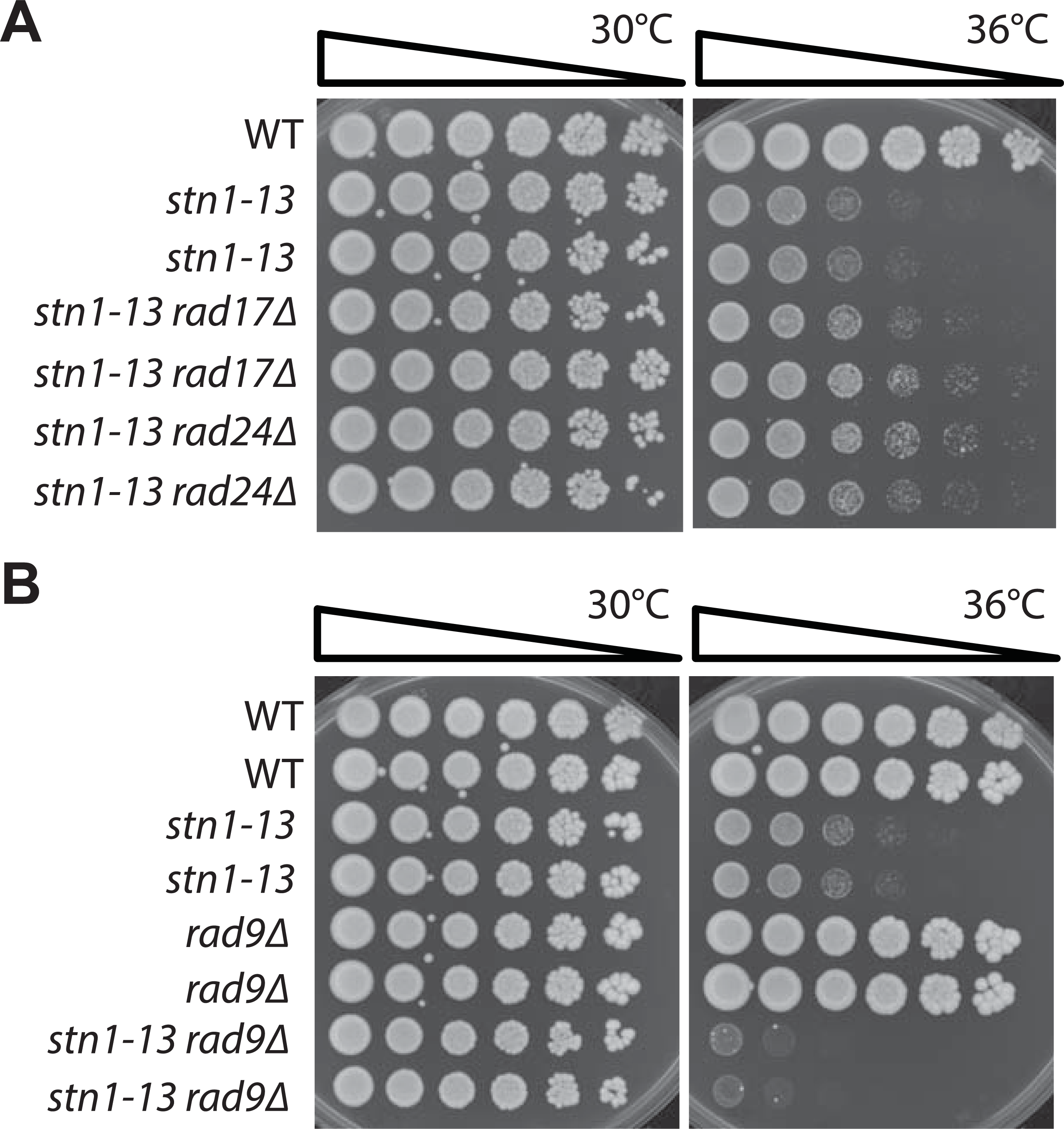
Effects of checkpoint and nonsense mediated decay pathways on fitness of *rfa3-313* strains. A-C) Saturated cultures of the yeast strains indicated (see Table 1) were five-fold serially diluted in water, spotted onto YEPD agar plates and incubated at the indicated temperatures for two days before being photographed.

**Figure S2:**
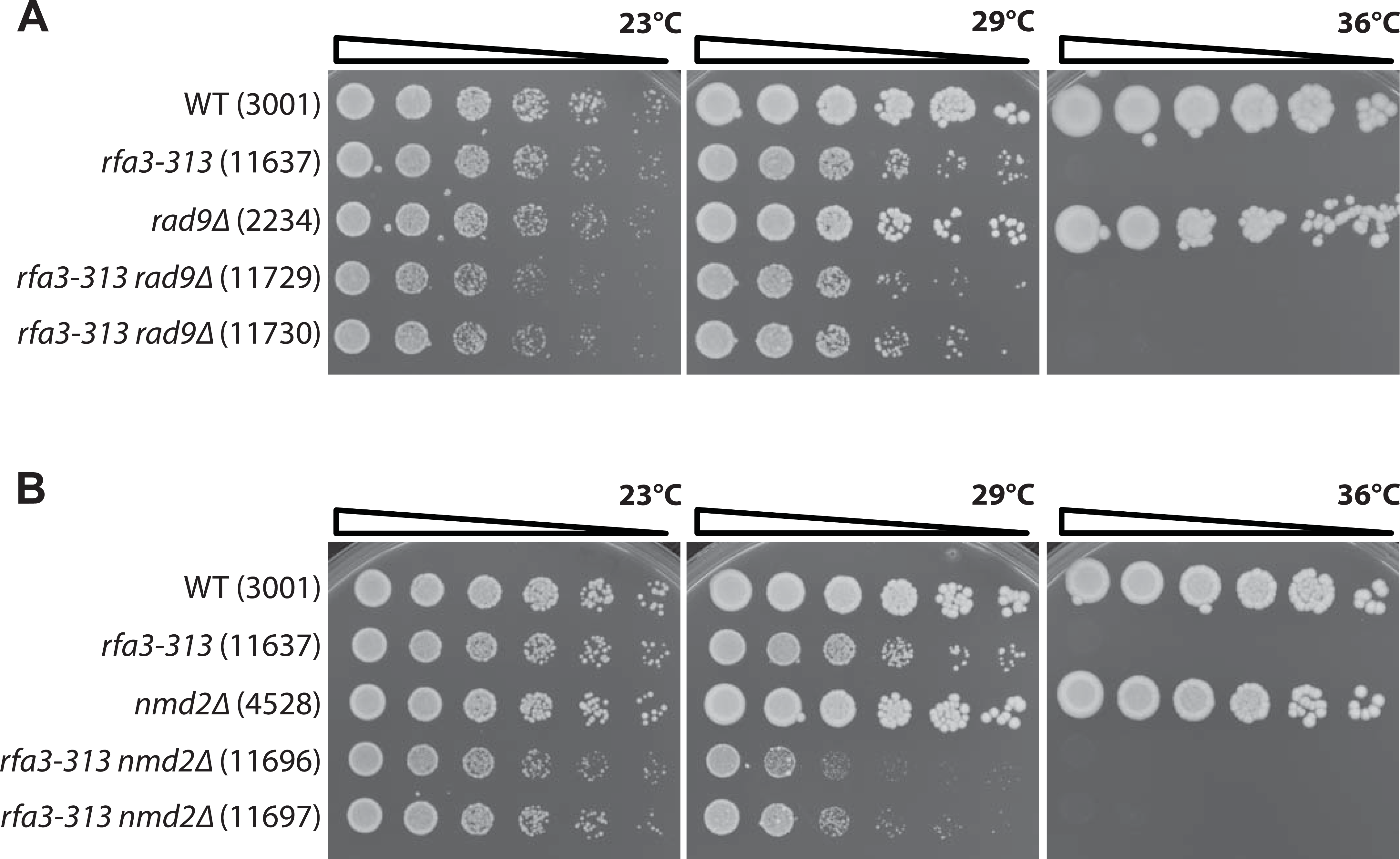
Effects of checkpoint pathways on fitness of *stn1-13* strains. A-B) Saturated cultures of the yeast strains indicated (see Table 1) were five-fold serially diluted in water, spotted onto YEPD agar plates and incubated at the indicated temperatures for two days before being photographed.

**Figure S3:**
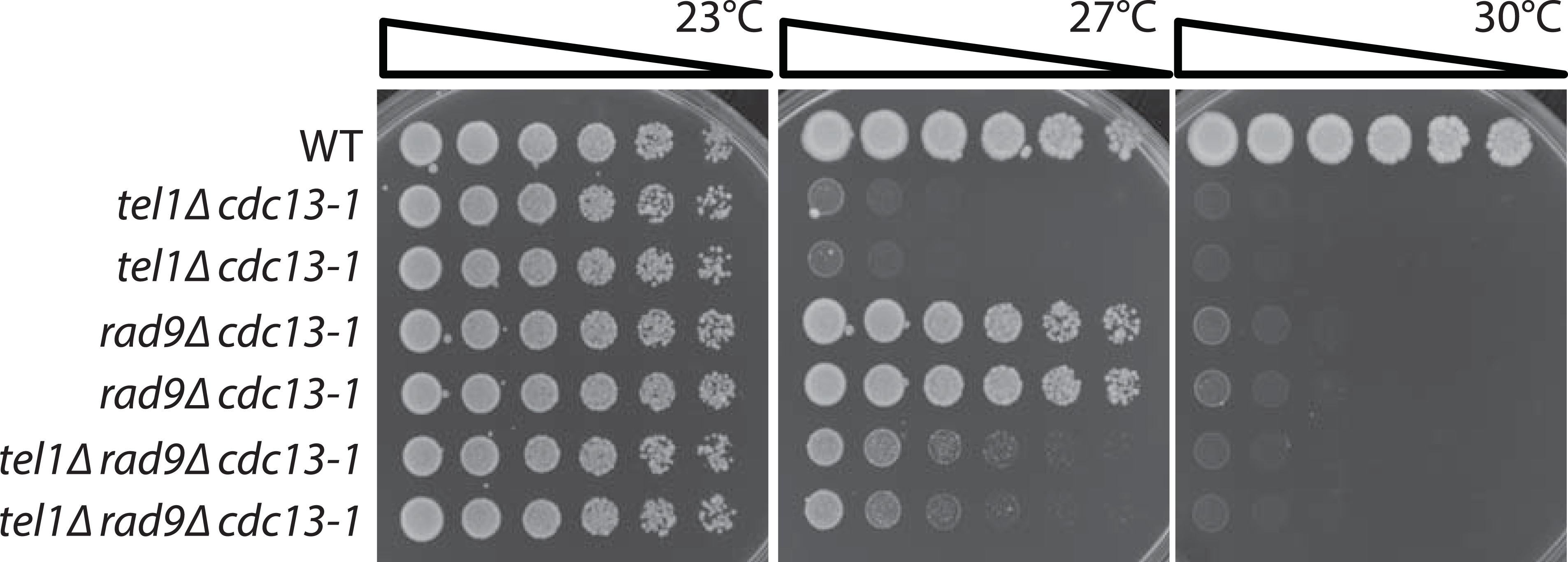
Effects of Tel1 and Rad9 on fitness of *cdc13-1* strains. Saturated cultures of the yeast strains indicated (see Table 1) were five-fold serially diluted in water, spotted onto YEPD agar plates and incubated at the indicated temperatures for two days before being photographed.

**Figure S4:**
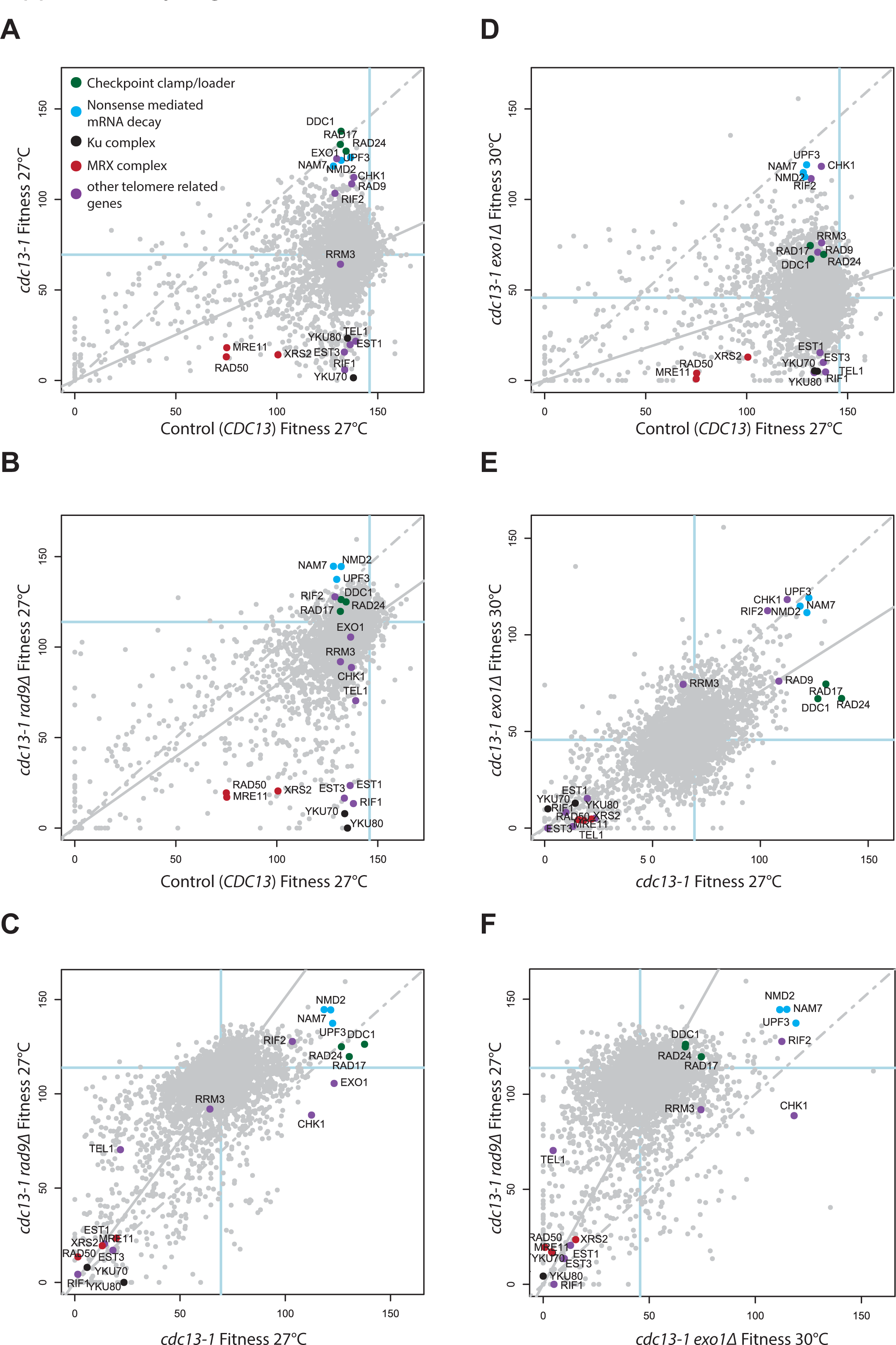
Examples of fitness plots and 2D scatter plots which can be generated with DIXY. A & C) Fitness plots, comparing fitness summaries for ˜5,000 *yfgΔ* deletions in a query screen that includes a background mutation (y-axis) and a matched control screen that differs from the query in that it lacks the background mutation (x-axis). Fitness plots include a prediction of double mutant fitness assuming genetic independence (solid line). Any deviations from this prediction are a measure of the strength of genetic interaction (see Addinall et al. (2011)). B, D, E, F) Regular 2D scatterplots generated by DIXY, which can also be a useful means to interrogate QFA data.

## References

Addinall, S. G., Downey, M., Yu, M., Zubko, M. K., Dewar, J., Leake, A., Hallinan, J., Shaw, O., James, K., Wilkinson, D. J., Wipat, A., Durocher, D. & Lydall, D. 2008. A genomewide suppressor and enhancer analysis of cdc13-1 reveals varied cellular processes influencing telomere capping in Saccharomyces cerevisiae. Genetics, 180, 180–2251.

Addinall, S. G., Holstein, E. M., Lawless, C., Yu, M., Chapman, K., Banks, A. P., Ngo, H. P., Maringele, L., Taschuk, M., Young, A., Ciesiolka, A., Lister, A. L., Wipat, A., Wilkinson, D. J. & Lydall, D. 2011. Quantitative fitness analysis shows that NMD proteins and many other protein complexes suppress or enhance distinct telomere cap defects. PLoS Genet, 7, e1001362.

Armanios, M. Y., Chen, J. J., Cogan, J. D., Alder, J. K., Ingersoll, R. G., Markin, C., Lawson, W. E., Xie, M., Vulto, I., Phillips, J. A., 3RD, Lansdorp, P. M., Greider, C. W. & Loyd, J. E. 2007. Telomerase mutations in families with idiopathic pulmonary fibrosis. N Engl J Med, 356, 356–1317.

Artandi, S. E. & Depinho, R. A. 2010. Telomeres and telomerase in cancer. Carcinogenesis, 31, 9–18.

Aubert, G. & Lansdorp, P. M. 2008. Telomeres and aging. Physiol Rev, 88, 88–557.

Bertuch, A. A. & Lundblad, V. 2006. The maintenance and masking of chromosome termini. Curr Opin Cell Biol, 18, 18–247.

Blackburn, E. H., Epel, E. S. & Lin, J. 2015. Human telomere biology: A contributory and interactive factor in aging, disease risks, and protection. Science, 350, 350–1193.

Booth, C., Griffith, E., Brady, G. & Lydall, D. 2001. Quantitative amplification of single-stranded DNA (QAOS) demonstrates that cdc13-1 mutants generate ssDNA in a telomere to centromere direction. Nucleic Acids Res, 29, 29–4414.

Chang, W., Cheng, J., Allaire, J. J., Xie, Y. & Mcpherson, J. 2015. shiny: Web Application Framework for R.. 0.12.2. ed.

Chen, L. Y., Redon, S. & Lingner, J. 2012. The human CST complex is a terminator of telomerase activity. Nature, 488, 488–540.

Dahlseid, J. N., Lew-SMITH, J., Lelivelt, M. J., Enomoto, S., Ford, A., Desruisseaux, M., Mcclellan, M., Lue, N., Culbertson, M. R. & Berman, J. 2003. mRNAs encoding telomerase components and regulators are controlled by UPF genes in Saccharomyces cerevisiae. Eukaryot Cell, 2, 2–134.

Dubarry, M., Lawless, C., Banks, A. P., Cockell, S. & Lydall, D. 2015. Genetic Networks Required to Coordinate Chromosome Replication by DNA Polymerases alpha, delta, and epsilon in Saccharomyces cerevisiae. G3 (Bethesda), 5, 5–2187.

Gao, H., Cervantes, R. B., Mandell, E. K., Otero, J. H. & Lundblad, V. 2007. RPA-like proteins mediate yeast telomere function. Nat Struct Mol Biol, 14, 14–208.

Garvik, B., Carson, M. & Hartwell, L. 1995. Single-stranded DNA arising at telomeres in cdc13 mutants may constitute a specific signal for the RAD9 checkpoint. Mol Cell Biol, 15, 15–6128.

Goulian, M., Heard, C. J. & Grimm, S. L. 1990. Purification and properties of an accessory protein for DNA polymerase alpha/primase. J Biol Chem, 265, 265–13221.

Gunes, C. & Rudolph, K. L. 2013. The role of telomeres in stem cells and cancer. Cell, 152, 152–390.

Holohan, B., Wright, W. E. & Shay, J. W. 2014. Cell biology of disease: Telomeropathies: an emerging spectrum disorder. J Cell Biol, 205, 205–289.

Holstein, E. M., Clark, K. R. & Lydall, D. 2014. Interplay between nonsense-mediated mRNA decay and DNA damage response pathways reveals that Stn1 and Ten1 are the key CST telomere-cap components. Cell Rep, 7, 7–1259.

Holstein, E. M. & Lydall, D. 2012. Quantitative amplification of single-stranded DNA. Methods Mol Biol, 920, 920–323.

Jia, X., Weinert, T. & Lydall, D. 2004. Mec1 and Rad53 inhibit formation of single-stranded DNA at telomeres of Saccharomyces cerevisiae cdc13-1 mutants. Genetics, 166, 166–753.

Lazzaro, F., Sapountzi, V., Granata, M., Pellicioli, A., Vaze, M., Haber, J. E., Plevani, P., Lydall, D. & Muzi-Falconi, M. 2008. Histone methyltransferase Dot1 and Rad9 inhibit single-stranded DNA accumulation at DSBs and uncapped telomeres. EMBO J, 27, 27–1502.

Lee, J. R., Xie, X., Yang, K., Zhang, J., Lee, S. Y. & Shippen, D. E. 2016. Dynamic interactions of Arabidopsis TEN1: stabilizing telomeres in response to heat stress. Plant Cell.

Luciano, P., Coulon, S., Faure, V., Corda, Y., Bos, J., Brill, S. J., Gilson, E., Simon, M. N. & Geli, V. 2012. RPA facilitates telomerase activity at chromosome ends in budding and fission yeasts. EMBO J, 31, 31–2034.

Lue, N. F., Chan, J., Wright, W. E. & Hurwitz, J. 2014. The CDC13-STN1-TEN1 complex stimulates Pol alpha activity by promoting RNA priming and primase-to-polymerase switch. Nat Commun, 5, 57–62.

Lydall, D. & Weinert, T. 1995. Yeast checkpoint genes in DNA damage processing: implications for repair and arrest. Science, 270, 270–1488.

Lydall, D. & Weinert, T. 1997. G2/M checkpoint genes of Saccharomyces cerevisiae: further evidence for roles in DNA replication and/or repair. Mol Gen Genet, 256, 256–638.

Maringele, L. & Lydall, D. 2002. EXO1-dependent single-stranded DNA at telomeres activates subsets of DNA damage and spindle checkpoint pathways in budding yeast yku70Delta mutants. Genes Dev, 16, 16–1919.

Markiewicz-POTOCZNY, M. & Lydall, D. 2016. Costs, benefits and redundant mechanisms of adaption to chronic low-dose stress in yeast. Cell Cycle, 15, 15–2732.

Morin, I., Ngo, H. P., Greenall, A., Zubko, M. K., Morrice, N. & Lydall, D. 2008. Checkpoint-dependent phosphorylation of Exo1 modulates the DNA damage response. EMBO J, 27, 27–2400.

Ngo, G. H., Balakrishnan, L., Dubarry, M., Campbell, J. L. & Lydall, D. 2014. The 9-1-1 checkpoint clamp stimulates DNA resection by Dna2-Sgs1 and Exo1. Nucleic Acids Res, 42, 42–10516.

Ngo, H. P. & Lydall, D. 2010. Survival and growth of yeast without telomere capping by Cdc13 in the absence of Sgs1, Exo1, and Rad9. PLoS Genet, 6, e1001072.

Petreaca, R. C., Chiu, H. C. & Nugent, C. I. 2007. The role of Stn1p in Saccharomyces cerevisiae telomere capping can be separated from its interaction with Cdc13p. Genetics, 177, 177–1459.

Polotnianka, R. M., Li, J. & Lustig, A. J. 1998. The yeast Ku heterodimer is essential for protection of the telomere against nucleolytic and recombinational activities. Curr Biol, 8, 8–831.

Price, C. M., Boltz, K. A., Chaiken, M. F., Stewart, J. A., Beilstein, M. A. & Shippen, D. E. 2010. Evolution of CST function in telomere maintenance. Cell Cycle, 9, 9–3157.

Sanchez, Y., Bachant, J., Wang, H., Hu, F., Liu, D., Tetzlaff, M. & Elledge, S. J. 1999. Control of the DNA damage checkpoint by chk1 and rad53 protein kinases through distinct mechanisms. Science, 286, 286–1166.

Sugitani, N. & Chazin, W. J. 2015. Characteristics and concepts of dynamic hub proteins in DNA processing machinery from studies of RPA. Prog Biophys Mol Biol, 117, 117–206.

Surovtseva, Y. V., Churikov, D., Boltz, K. A., Song, X., Lamb, J. C., Warrington, R., Leehy, K., Heacock, M., Price, C. M. & Shippen, D. E. 2009. Conserved telomere maintenance component 1 interacts with STN1 and maintains chromosome ends in higher eukaryotes. Mol Cell, 36, 36–207.

Tong, A. H. & Boone, C. 2006. Synthetic genetic array analysis in Saccharomyces cerevisiae. Methods Mol Biol, 313, 313–171.

Tong, A. H., Evangelista, M., Parsons, A. B., Xu, H., Bader, G. D., Page, N., Robinson, M., Raghibizadeh, S., Hogue, C. W., Bussey, H., Andrews, B., Tyers, M. & Boone, C. 2001. Systematic genetic analysis with ordered arrays of yeast deletion mutants. Science, 294, 294–2364.

Wellinger, R. J. & Zakian, V. A. 2012. Everything you ever wanted to know about Saccharomyces cerevisiae telomeres: beginning to end. Genetics, 191, 191–1073.

Zubko, M. K., Guillard, S. & Lydall, D. 2004. Exo1 and Rad24 differentially regulate generation of ssDNA at telomeres of Saccharomyces cerevisiae cdc13-1 mutants. Genetics, 168, 15–103.

